# A comprehensive analysis of the potential biological functions and prognostic values of SREBF1 for multiple cancer types including colorectal cancer

**DOI:** 10.1101/2024.07.26.605404

**Authors:** Dongling Li, Fan Xu, Qinrui Cai, Ling Lin, Xiaoya Zhou, Li Li, Yao Chen, Tianlin Feng, Yuanxiu Gan, Chenhua Zhang, Fan Yang

## Abstract

As an important molecule involved in lipid metabolism, Sterol Regulatory Element-Binding Protein 1 (SREBF1) plays a critical role in governing cellular lipid synthesis and uptake. However, a comprehensive understanding of the biological significance of SREBF1 in pan-cancer remains elusive. In this study, we aimed to comprehensively analyze the functional characteristics of SREBF1 in human cancers, including colorectal cancer (CRC). Our study utilized various databases and tools such as TCGA, GEO, TIMER2.0, GEPIA2.0, UALCAN, and the Human Protein Atlas to examine SREBF1 expression, genetic alterations, immune infiltration, single-cell sequencing, survival analysis, and gene enrichment. The results revealed that the expression of SREBF1 exhibited significant variations across different cancer types, and was closely associated with patient prognosis and immune infiltration. Moreover, the gene function assays indicated SREBF1 significantly affect cell growth and migration. Overall, the findings suggest that SREBF1 could serve as a potential prognostic biomarker and a novel therapeutic target in different cancer types.

## Introduction

Cancer is a disease marked by the abnormal growth of cells, which often spreading to nearby tissues. It can occur in any part of the body due to genetic factors, environmental influences, or lifestyle choices[1, 2]. And pan-cancer analysis reveals commonalities and variations in human malignancies by studying molecular abnormalities across different cancer types[3]. Utilizing resources like TCGA and GEO, this analysis aids in developing combination and personalized therapies for diverse cancer models[4, 5].

Sterol Regulatory Element-Binding Transcription Factor 1 (SREBF1) is a specific isoform of the Sterol Regulatory Element-Binding Transcription Factor (SREBF) family[6]. It is also known as SREBP-1 or SREBP-1a. SREBF1 plays a crucial role in regulating the expression of genes involved in the biosynthesis of fatty acids, triglycerides, and cholesterol. It is highly expressed in tissues involved in regulating of lipid metabolism, including liver, adipose tissue, and muscle[7, 8].

SREBF1 is activated in response to low cellular levels of lipids, which leads to the upregulation of genes involved in lipogenesis and lipid uptake. This process is critical for maintaining energy homeostasis and preventing metabolic disorders. Additionally,SREBF1 has been linked a number of other biological processes, including inflammation, cell proliferation, and differentiation.The dysregulation of SREBF1 activity has been associated with a number of pathological conditions, including cardiovascular disease, certain types of cancer, and non-alcoholic fatty liver disease.[9, 10]. Studies have suggested that SREBF1 may have a role in regulating immune function. One of how SREBF1 could modulate immune function is through its regulation of lipid metabolism, as lipids and cholesterol play an important role in immune cell function. But the precise role of SREBF1 in immune function is still being elucidated.

This research provides a comprehensive analysis of SREBF1 expression across multiple cancer types. The analysis encompasses differential gene expression, protein correlation, pathway, and prognostic analysis of various tumor types and stages. Additionally, the study sought to ascertain the association between SREBF1 expression and the presence of immune infiltrating cells. Meanwhile, gene enrichment analysis highlighted the involvement of SREBF1-associated molecules in tumorigenesis. All the findings indicated that SREBF1 could be a valuable prognostic biomarker in many types of cancers as a novel therapeutic strategy.

### Materials and methods Gene expression analysis

The TIMER2.0 database (http://timer.cistrome.org/) was used to examine the expression level of SREBF1 between the pan-cancerous tissue and the adjacent normal tissue.[11]. The GEPIA2.0 database(http://gepia2.cancer-pku.cn/#index)was used to assess the association between patients’ pathological stage and SREBF1 expression in all TCGA cancers[12]. The CPTAC within the UALCAN database (http://ualcan.path.uab.edu/index.html) was utilized to examine the protein expression of SREBF1 across various types of cancers[13, 14]. The Human Protein Atlas (HPA) database (https://www.proteinatlas.org) was available to study the expression levels of SREBF1 between the tumor and its corresponding tissues[15]. The String database was used to build the protein-protein interaction network (PPI) for SREBF1[16]. The GEPIA 2.0 database (http://gepia2.cancerpku.cn/#index) could analyze the correlation between SREBF1 and other genes in multiple cancers of TCGA data.

### Genetic alteration analysis

The cBioPortal database(https://www.cbioportal.org/)was utilized to gather the data about the alteration frequency, mutation type, mutation site information, and threedimensional (3D) structure of SREBF1 in all TCGA databases[17]. The promoter methylation levels of SREBF1 in various types of cancers were analyzed by the UALCAN database (http://ualcan.path.uab.edu/index.html).

### The infiltration of immune cells

We analyzed the association between SREBF1 and the immune infiltration levels in different cancer types by using the TIMER2.0 (http://timer.cistrome.org/), and we used Pearson’s correlation coefficient to calculate the correlation between the SREBF1 and various immune cell types, including CD4+ T cells, CD8+ T cells, neutrophils, macrophages, eosinophils, and natural killer cells.

### Single-cell sequencing

CancerSEA database (http://biocc.hrbmu.edu.cn/CancerSEA/) as a professional single cell sequencing, we utilized it to analyze the relationship between the single-cell level and SREBF1’s expression, then generated the results of RB, UM, and AML by T-SNE diagram[17]. We used the Singerbox (http://www.sangerbox.com/) to obtain the heatmap between the SREBF1 and the immune correlation.

### Survival analysis

Using the GEPIA2 analysis to assess the relationship between the expression of SREBF1 and the prognostic of patients, including overall survival (OS), progression-free survival (PFS), disease-free survival (DFS), post-progression survival (PPS), decreased event-free survival (EFS), and failure point (FP), and the patients divided into the low expression and high expression groups. The hazard ratio was calculated based on the Cox PH Model.

### Gene enrichment analysis

The GEPIA2.0 database(http://gepia2.cancer-pku.cn/#index)included the TCGA and GTEx data, which could generate the top 100 SREBF1-related genes, and then we conducted the heatmap between the top six related genes and SREBF1. Gene ontology (GO) and Kyoto Encyclopedia of Genes and Genomes (KEGG) enrichment analyses were employed to analyze the SREBF1-influenced biological functions and signaling pathways in TCGA tumors. P-value < 0.05 was thought to be statistically significant.

### Cell lines

SW480, HCT116, NCM460, LS174T, HT29, and LOVO were bought from Cell Resource Center, Peking Union Medical College (PCRC).

### Quantitative PCR (qPCR)

Total RNA was extracted using Steady Quick RNA Extraction Kit (ACCURATE BIOLOGY), then used ABScript III RT Master Mix for qPCR with gDNA Remover (ABclonal) reverse transcription into cDNA by following the protocol. RT-qPCR was conducted using 2X Universal SYBR Green Fast qPCR Mix (ABclonal), according to the manufacturer’s instructions. Beta-Actin was used to normalize the expression of the target genes. Primer sequences for RT-qPCR were as follows: primer for SREBF1 (forward primer: CGGAACCATCTTGGCAACAGT, reverse primer: CGCTTCTCAATGGCGTTGT) and primer for Beta-Actin (forward primer: CATGTACGTTGCTATCCAGGC, reverse primer: CTCCTTAATGTCACGCACGAT).

### Western blot

The harvested cells were washed with the 1X iced-PBS and lysed in RIPA cell lysis buffer for 5 minutes, then ultrasonic the mixed buffer for 10 min, and centrifugation at 14,000 × g, 4 °C for 15 min, the supernatant was protein for us to analysis. The protein lysates were separated on SDS-PAGE gels and transferred onto a 0.45mm PVDF membrane for 2 hours. The membrane was then subjected to overnight incubation at 4 °C with SREBF1 antibodies (Proteintech). The next day, after three washes in TBST, the membrane was incubated with the appropriate secondary antibody for 1 hour at room temperature. Finally, the membrane was exposed by the VILBER image-forming system.

### Cell Growth Assay

A total of 2000 cells were seeded in 96-well plates and cultured for 5 days. Thereafter, 10 μL of CCK-8 solution was added to each well daily. After incubation for 60 minutes at 37°C, the optical density (OD) values at 450 nm were determined using a microplate reader (Tecan, Austria) and normalised to the corresponding control.

### Colony formation Assay

In total, 1000 cells were plated in 6-well plates and cultured for approximately 14 days. Cell colonies were fixed with 4% formaldehyde (P0099, Beyotime) and stained with 0.1% crystal violet (C8470, Solarbio) for 15 minutes, and the colonies were photographed and counted manually.

### Transwell Assay

Migration assays were performed using Transwell chambers in the presence of Matrigel (Corning 354,480). SREBF1 knockdown HCT116 cells (1 × 10^5^ cells/well) were seeded in the upper chamber with serum-free medium, and 700 µl of 10% FBS was added to the lower chamber of the 24-well plate. After 48 hours of incubation, the cells on the upper surface of the filter were completely removed and washed by PBS. The filters were then fixed in 4% paraformaldehyde and stained with crystal violet (Beyotime C0121).

## Results

### 1. The expression of SREBF1 in human pan-cancer

To better understand the effect of SREBF1, we initially evaluated the level of SREBF1 expression in different cancers using TIMER2.0. As Figure. 1A shows, compared the expression between tumor and adjacent non-tumor tissues, SREBF1 expression was significantly up-regulated in Bladder Urothelial Carcinoma(BLCA), Breast Cancer (BRCA), Cervical Squamous Cell Carcinoma (CESC), Colon Adenocarcinoma (COAD), Esophageal Squamous Cell Carcinoma (ESCA), Glioblastoma (GBM), Head and Neck Squamous Cell Carcinoma (HNSC), HPV-Positive Head and Neck Squamous Cell Carcinoma (HNSC-HPV+) and HPV-Negative Head and Neck Squamous Cell Carcinoma (HNSC-HPV-), Kidney Chromophobe (KICH), Kidney Renal Clear Cell Carcinoma (KIRC), Kidney Renal Papillary Cell Carcinoma (KIRP), Lung Adenocarcinoma (LUAD), Lung Squamous Cell Carcinoma (LUSC), Prostate Adenocarcinoma (PRAD), Skin Cutaneous Melanoma (SKCM), Stomach Adenocarcinoma (STAD), Thyroid Carcinoma (THCA), Uterine Corpus Endometrial Carcinoma (UCEC) and so on. Only in GBM and Pheochromocytoma and Paraganglioma (PCPG), adjacent normal tissues have higher expression of SREBF1. Next, based on these data, our further analysis revealed that the expression level of SREBF1 in Pediatric Brain Cancer varies greatly with age. And the SREBF1 with less expression in Pediatric Brain Cancer recurrence or progression than the initial tumor with the UALCAN database (Supplementary Figure.s1).

**Figure.1.**
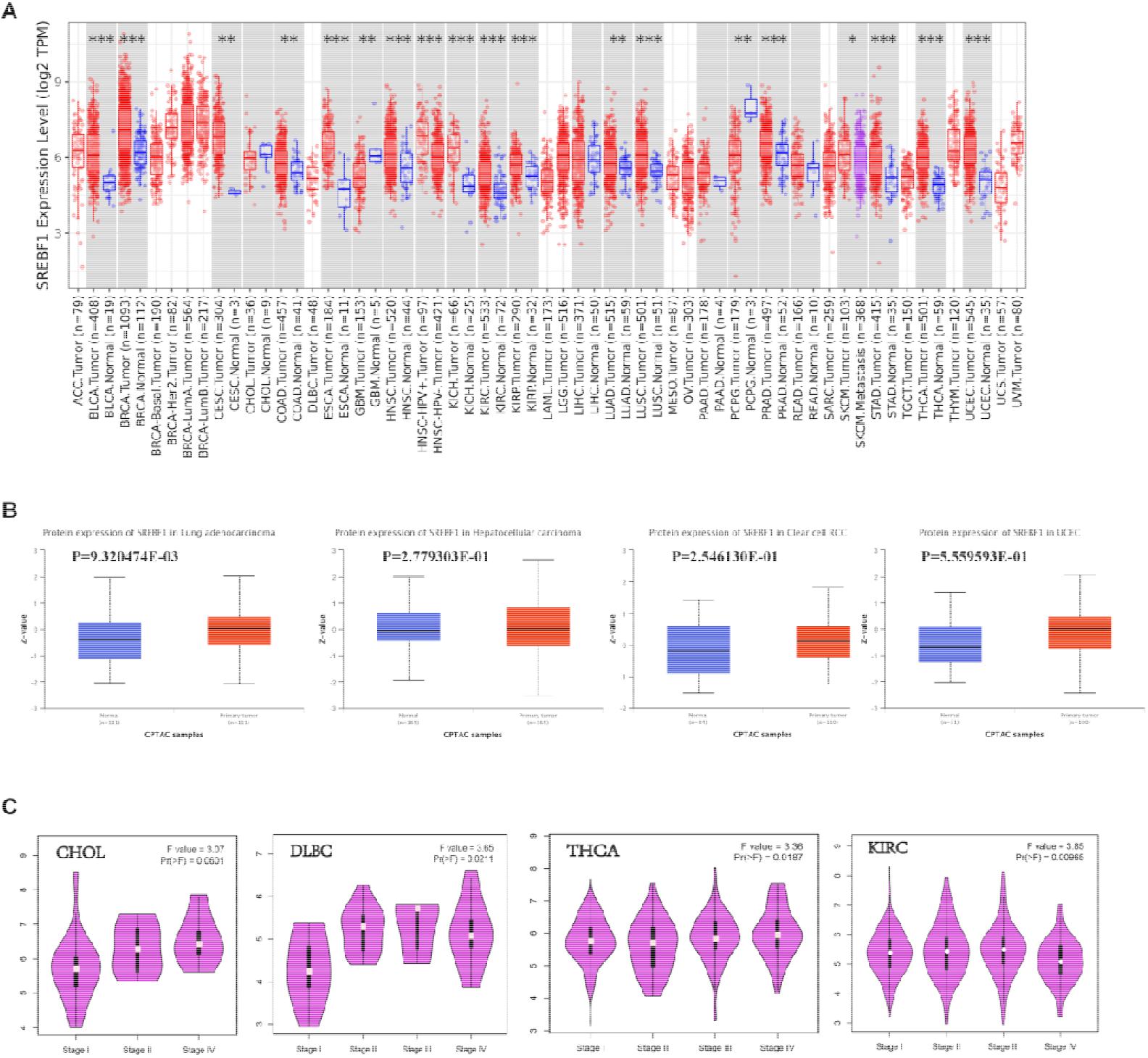
The expression of SREBF1 in pan-cancer. (A) SREBF1 expression in different cancers from TIMER2.0. *p < 0.05; **p < 0.01; ***p < 0.001. (B) The protein levels of SREBF1 in LUAD, HCC, RCC and UCEC were analyzed using CPTAC. (C) SREBF1 expression levels and the pathological stages were analyzed using GEPIA2.0. (D)RT-PCR result of SREBF1 expression in human colorectal normal and tumor cell lines, p < 0.01. (E) Western blotting of SREBF1 differential expression in human colorectal normal and tumor cell lines.

Then, the protein levels of SREBF1 in pan-cancer tissues were further analyzed through the UALCAN database. The follow ing results demonstrated that the expression levels of the total SREBF1 protein were elevated in primary tissues of endometrioid cancer (EC), clear cell renal cell carcinoma (ccRCC), hepatocellular carcinoma (HCC), and LUAD compared with normal tissues (Figure. 1B). Besides, by utilizing GEPIA2.0, we explored the connection between SREBF1 expression levels and tumor stages, which led us to discover a significant effect of SREBF1 expression on the stages of the patients with CHOL, Diffuse Large B-Cell Lymphoma (DLBC), KIRC, and THCA (Figure. 1E).

Additionally, we verified the expression of SREBF1 by analyzing the IHC results sourced from the HPA database. Our analysis suggested that SREBF1 was predominantly strongly or positively expressed in tumor tissue originating from lung, prostate, breast, and melanoma (Figure. 2).

**Figure.2.**
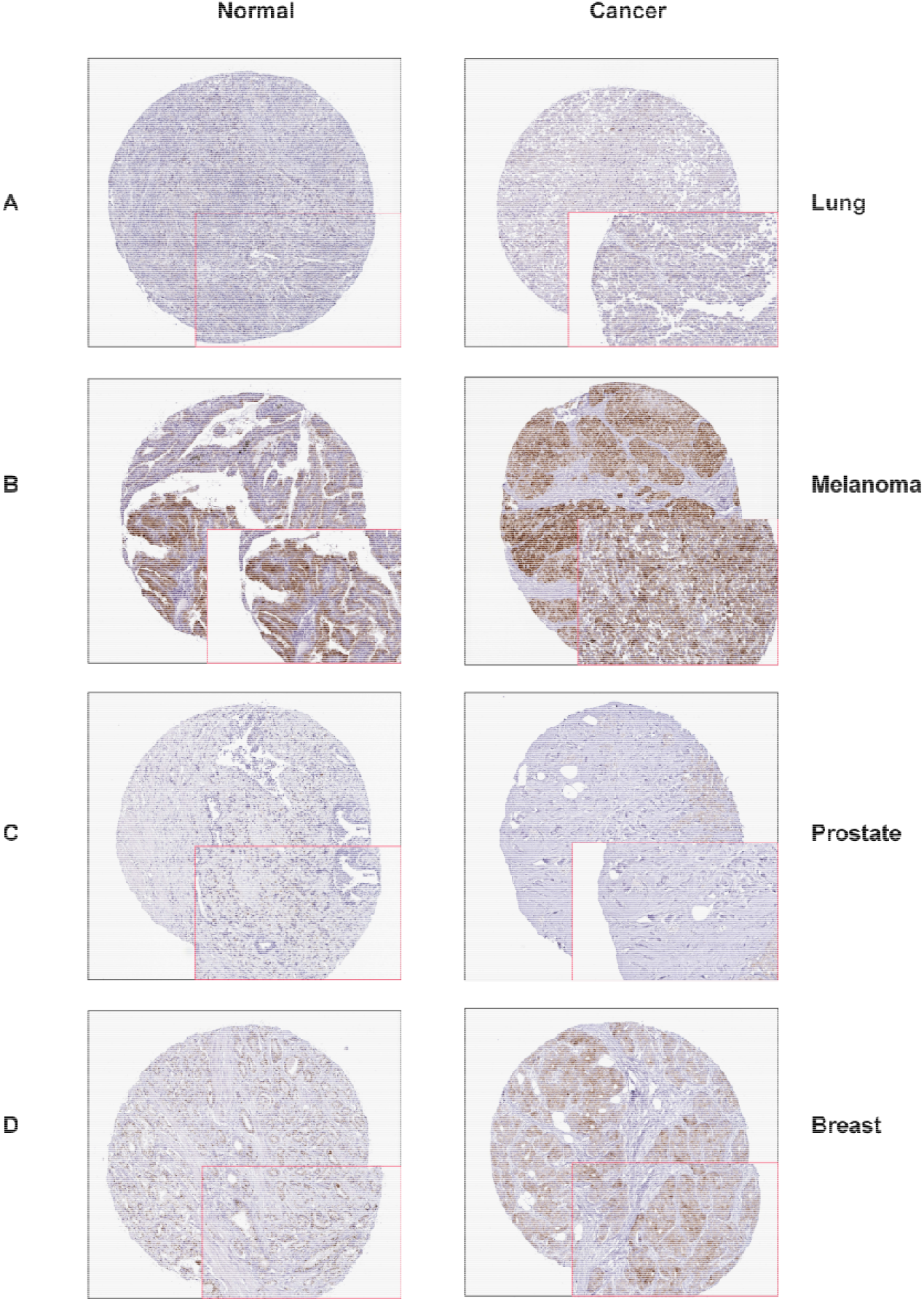
The different expression of SREBF1 between normal tissues and tumor tissues. (A–D) HPA platforms displayed the upregulated expression of SREBF1 in tumor tissue derived from lung(A, melanoma(B,prostate(C and breast(D.

### 2. Prognostic Assessment Value of SREBF1 in Pan-Cancer

We initially assessed the prognostic value of SREBF1 in pan-cancer OS. SREBF1 emerged as a risk factor in three tumor types, it also exhibited a favorable protective effect in several cancer types (Figure. 3A). Next, we used the GEPIA2.0 database to assess the prognostic value of SREBF1 in pan-cancer OS and DFS. We found that high expression of SREBF1 associated with poor OS in BLCA (p=0.016), Mesothelioma (MESO) (p=0.041), Acute Myeloid Leukemia (LAML) (p=0.011), and DFS analysis data indicated the correlation between high SREBF1 expression and poor DFS for cancers of BRCA (p=0.018), COAD (p=0.029), KIRC (0.015), and PCPG (p=0.029) (Figure. 3B/3C). Overall, the findings imply that SREBF1 has the ability to serve as a prognostic marker in multiple types of cancer.

**Figure.3.**
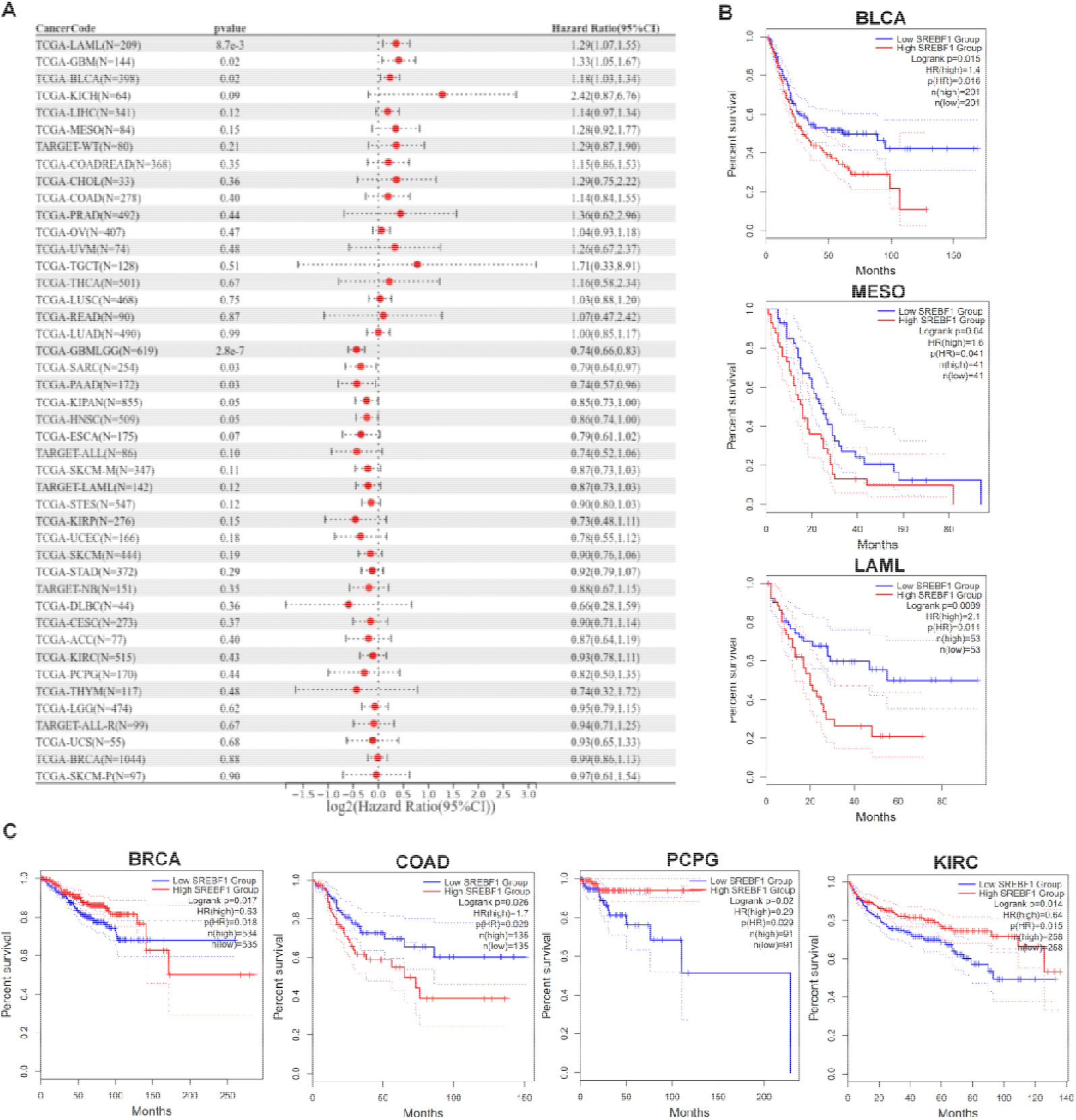
Prognostic values of SREBF1 expression in pan-cancer. (A) Forest plot of the correlation between SREBF1 and overall survival (OS) in patients with tumors.We used the GEPIA2 tool to investigate the impact of SREBF1 gene expression on patients’ prognosis, including overall survival (OS, B) and disease-free survival (DFS, C).

### 3. Investigation of genetic alterations of SREBF1 in Pan-cancer

By utilizing the cBioPortal database relay on TCGA data, we explored the gene mutations of SREBF1 in various cancers. As shown in Figure.4A, the most common DNA change was mutation. Patients with sarcoma (SAR), UCEC, STAD, SKCM, ESCA, Liver Hepatocellular Carcinosarcoma (LHC), BLCA, COAD, CESC, MESO, LUAD, LUSC, HNSC, PRAD, KIRP, ovarian serous cystadenocarcinoma, BRCA, AML, Brain Lower Grade Glioma (LGG), GBM, and KIRC showed SREBF1 mutation. Additionally, the analysis results showed the high SREBF1 amplification in SARC and UCS (>2%) (Figure. 4A). The sites, types, and case numbers of the SREBF1 gene modification were further displayed, showed that the misssence was the main mutation type of SREBF1 (Figure. 4B). We also observed the R586H/C site in the 3D model of the SREBF1 protein (Figure. 4C).

**Figure.4.**
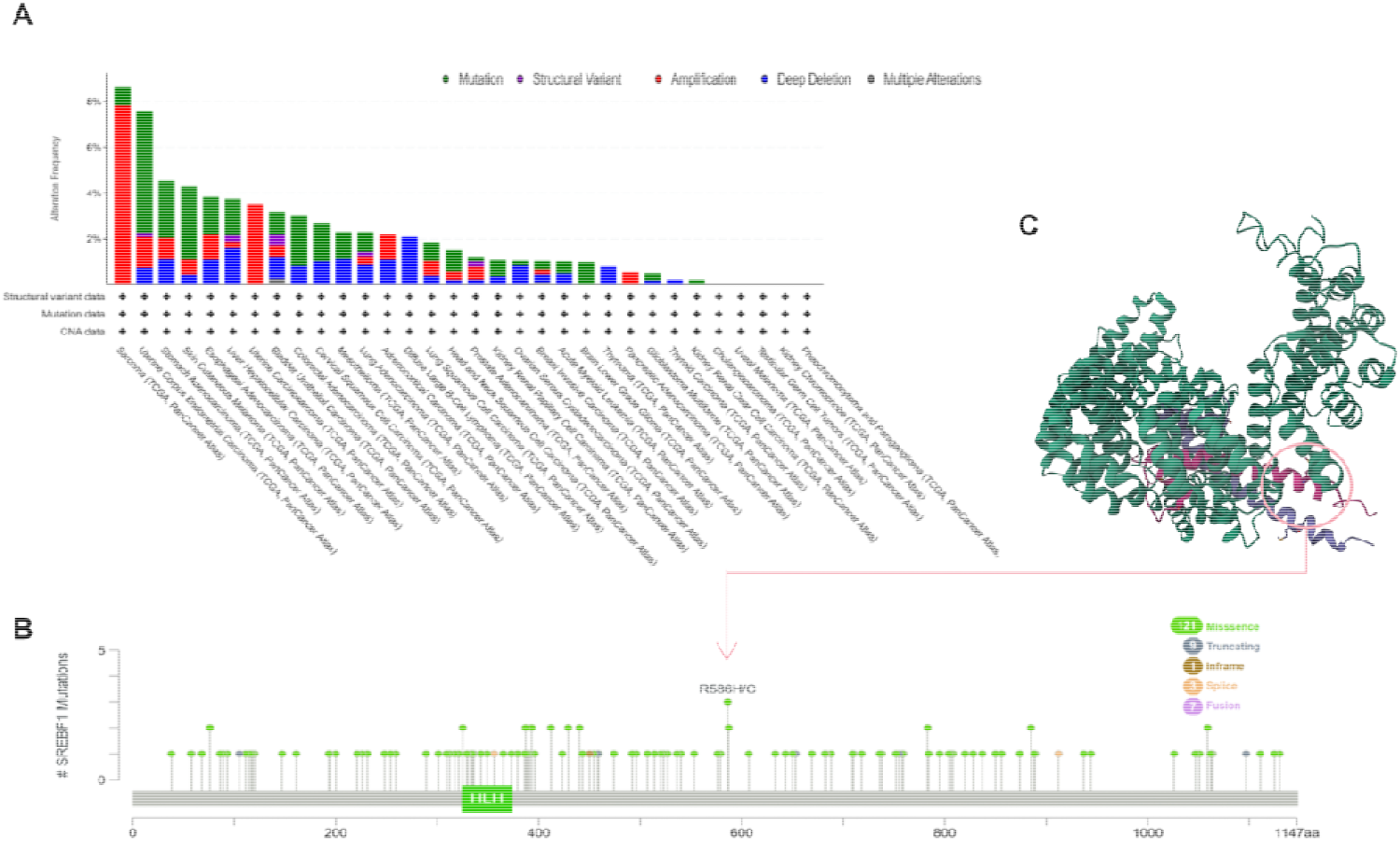
SREBF1 gene mutation in various cancers. (A,B) cBioPortal was used to display the alteration frequency of different mutation types (A) and mutation site (B) of SREBF1 in pan-cancer. (C) SREBF1 mutation site was shown in the 3D protein structure of SREBF1.

DNA methylation is closely related to cancer occurrence and progression[18, 19]. We investigated the DNA methylation of SREBF1 by the UALCAN. A marked decrease of SREBF1methylation level was observed in BLCA, THCA, CESC, LIHC, COAD, KIRC, PCPG, UCEC, and PAAD tissues compared to normal tissues by the UALCAN database. The methylation level of SREBF1 in HNSC, GBM, ESCA, BRCA, KIRP, THYM, CESC, LUSC, STAD, CHOL, PRAD, and SARC was increased (Figure. 5).

**Figure.5.**
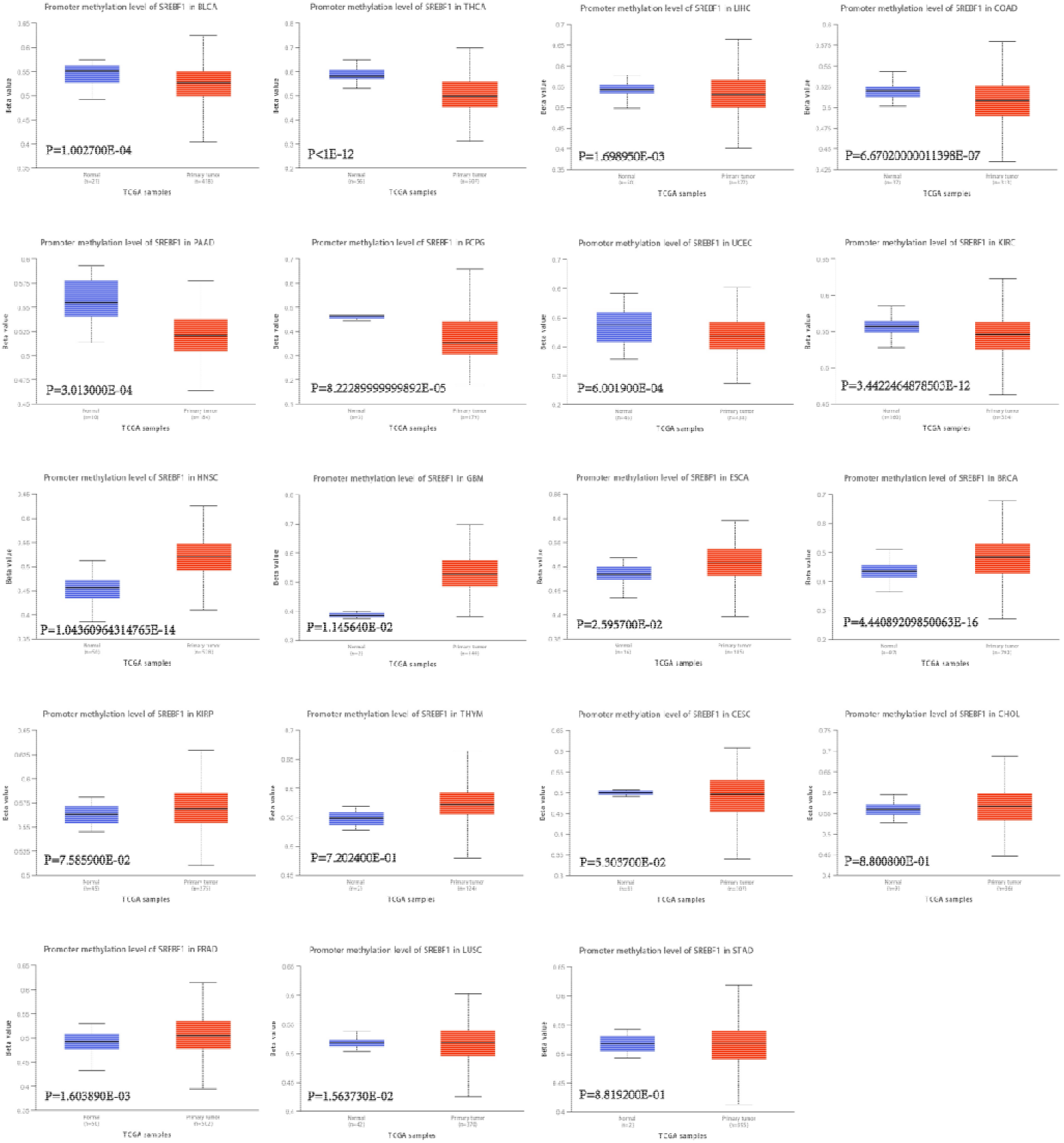
Promoter methylation levels of SREBF1 in cancers. The methylation values of SREBF1 between normal and primary tumor tissues were analyzed using UALCAN tool.

### 4. Correlation between the Immune Infiltration Cells and SREBF1

Immune cells play an important role in the tumor microenvironment (TME), and plenty of evidence indicated that cancer cells could interact with various components of the TME, then promote immune evasion and ultimately drive tumor growth, recurrence, and metastasis[20, 21]. To identify new targets and biomarkers of the cancer which can help develop more targeted and personalized immunotherapies that can effectively attack cancer cells while minimizing harm to healthy cells.

Our investigation found a positive relationship between the infiltration of CD8+ T cells and the expression of SREBF1 in DLBC and HNSC-HPV+. Similarly, the SREBF1 in HNSC, HNSC-HPV+, HNSC-HPV-, KICH, LIHC, LUAD, Ovarian Cancer (OV), and THCA has the same correlation with the infiltration of Tregs (Figure. 6A/6B). Furthermore, there was a significantly negative correlation between SREBF1 and the myeloid dendritic cells (mDCs) infiltration in CHOL (Figure. 6C). Our analysis didn’t find an obvious correlation between SREBF1 and the infiltration of natural killer cells (NK), neutrophils, CD4+ T cells, B cells, cancer-associated fibroblasts, and macrophages (Supplementary Figure.s2). Immune checkpoints play a crucial role as key constituents in the clinical immunotherapy for tumors. The association between the SREBF1 expression and checkpoints in pan-cancer was also evaluated. And the finding demonstrated a strong correlation between elevated levels of immune checkpoints and the SREBF1 in ACC, notably KIR2DL3, IL2RA, CD80, IFNG, BTN3A2, and IL1A (Figure. 6D).

**Figure.6.**
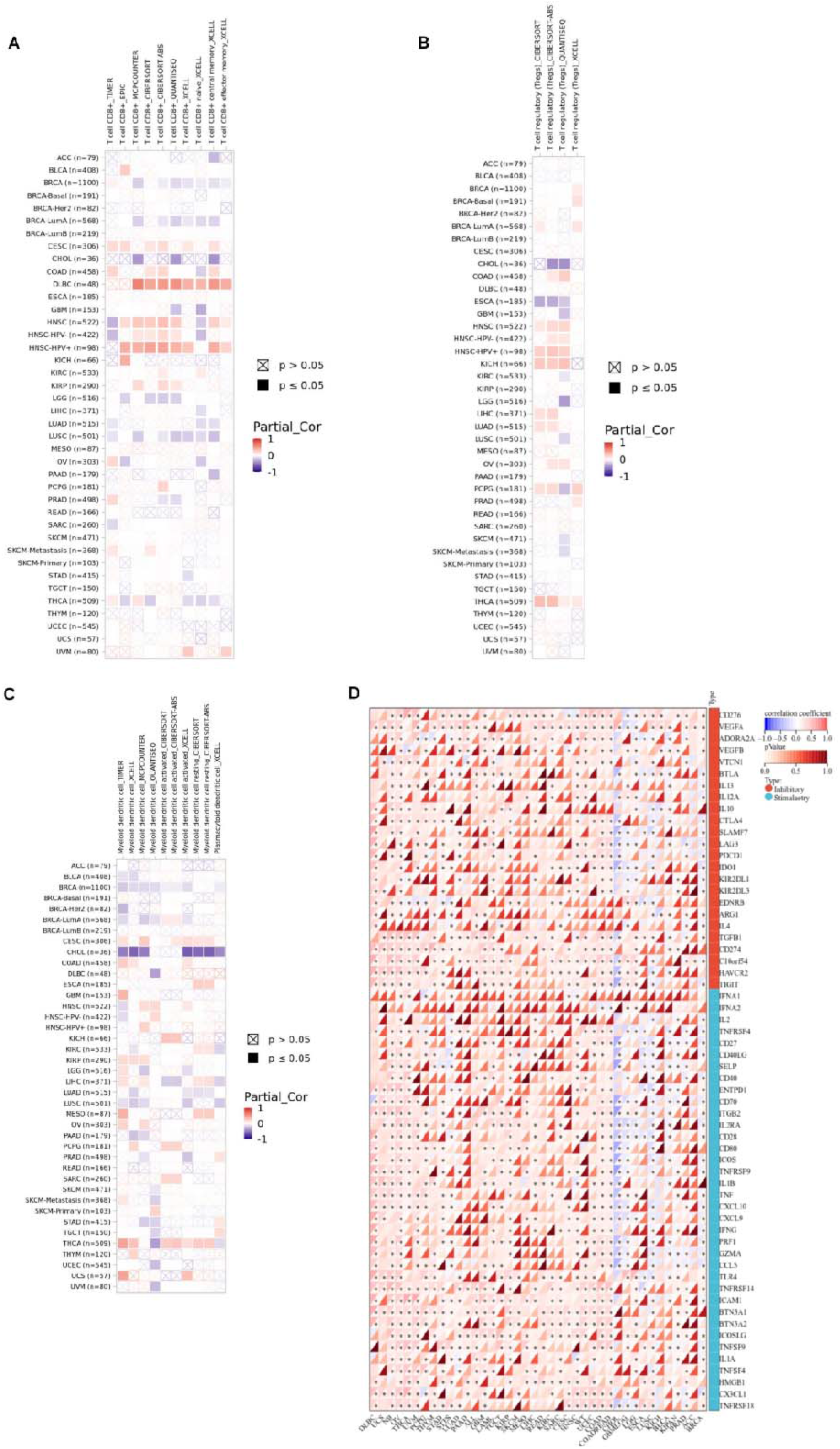
The correlation between immune cells and SREBF1 expression in cancers. (A–C) The relationship between SREBF1 expression and immune infiltration of CD8+ T cell (A), Tregs (B) and mDCs(C) was depicted by TIMER2.0 database. D.Correlation analyses of the SREBF1 expression with immune checkpoint genes in pan-cancer. *p < 0:05, **p < 0:01, ***p < 0:001, and ****p < 0:0001.Positive correlation(0–1) are indicated with the red color, while negative correlation (−1 to 0) are indicated with the blue color. p-value < 0.05 is considered as statistically significant. A cross indicates non-significant correlations.

### 5. The expression of SREBF1 at single-cell levels

Single-cell transcriptome sequencing is a molecular biology technique that allows for the analysis of gene expression at the level of individual cells. SREBF1 expression in RB was found to be negatively correlated with angiogenesis and differentiation (Figure. 7B). Also, SREBF1 expression was positively related with cell cycle, DNA repair, and DNA damage. In uveal melanoma (UM), SREBF1 expression was positively related to most tumor biological behaviors such as DNA repair, DNA damage, invasion, apoptosis, and metastasis (Figure. 7A). In addition, SREBF1 expression profiles were shown at single-cell levels from RB, ODG, and HNSCC through the T-SNE diagram (Figure. 7C).

**Figure.7.**
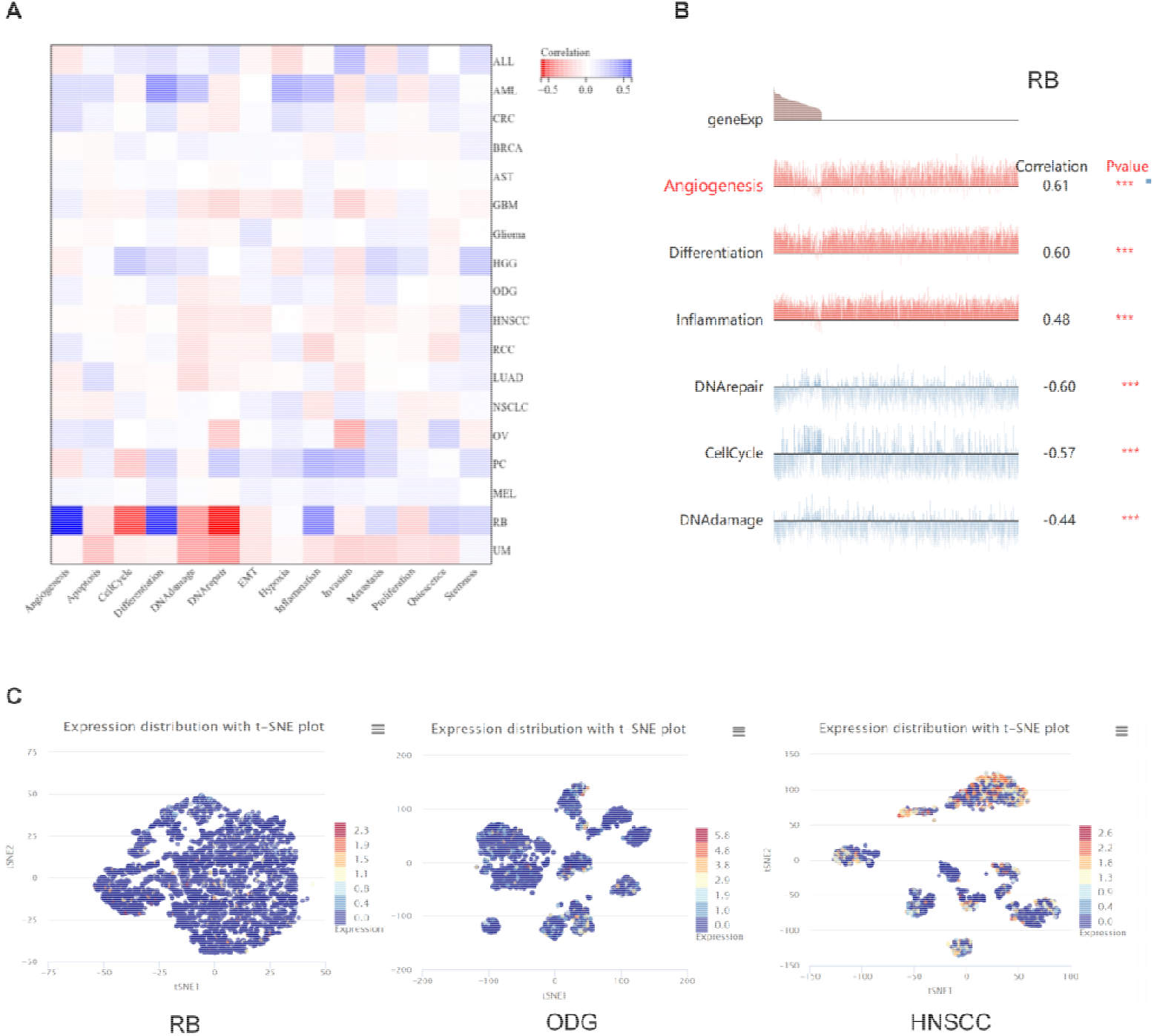
The expression levels of SREBF1 at single-cell levels. (A,B) The relationship between SREBF1 expression and different functional states in tumors was explored by the CancerSEA tool. *p < 0.05; **p < 0.01; ***p < 0.001. (C) SREBF1 expression profiles were shown at single cells from RB, ODG and AML HNSCC diagram.

### 6. Functional enrichment analysis of SREBF1 in cancers

To gain deeper insights into the molecular mechanism of the SREBF1 gene in tumors, we conducted a functional enrichment analysis on SREBF1-associated binding proteins as well as genes that exhibit expression correlation with SREBF1. By utilizing the STRING tool, we identify 10 binding proteins that have been experimentally validated to interact with SREBF1. Subsequently, a protein-protein interaction (PPI) network was then constructed for these proteins to further elucidate their relationship with SREBF1 (Figure. 8A). We also by utilizing the GEPIA2.0 tool obtained the top 100 genes associated with SREBF1 expression in pan-cancer. Among these genes, X-box binding protein 1 (XBP1), potassium channel tetramerization domain containing 3 (KCTD3), rabaptin, RAB GTPase binding effector protein 1 (RABEP1), transcriptional repressor GATA binding 1 (TRPS1), GATA binding protein 3 (GATA3), LIM homeobox transcription factor 1 beta (LMX1B) observed strong interaction s with SREBF1 in the vast of cancer types. (Figure. 8B/8C). We also performed enrichment analysis using KEGG and GO, and the results indicated that SREBF1 might potentially participate in multiple pathways, including prostate gland development, signaling by nuclear receptors, and omega-9 fatty acid synthesis (Figure. 8D).

**Figure.8.**
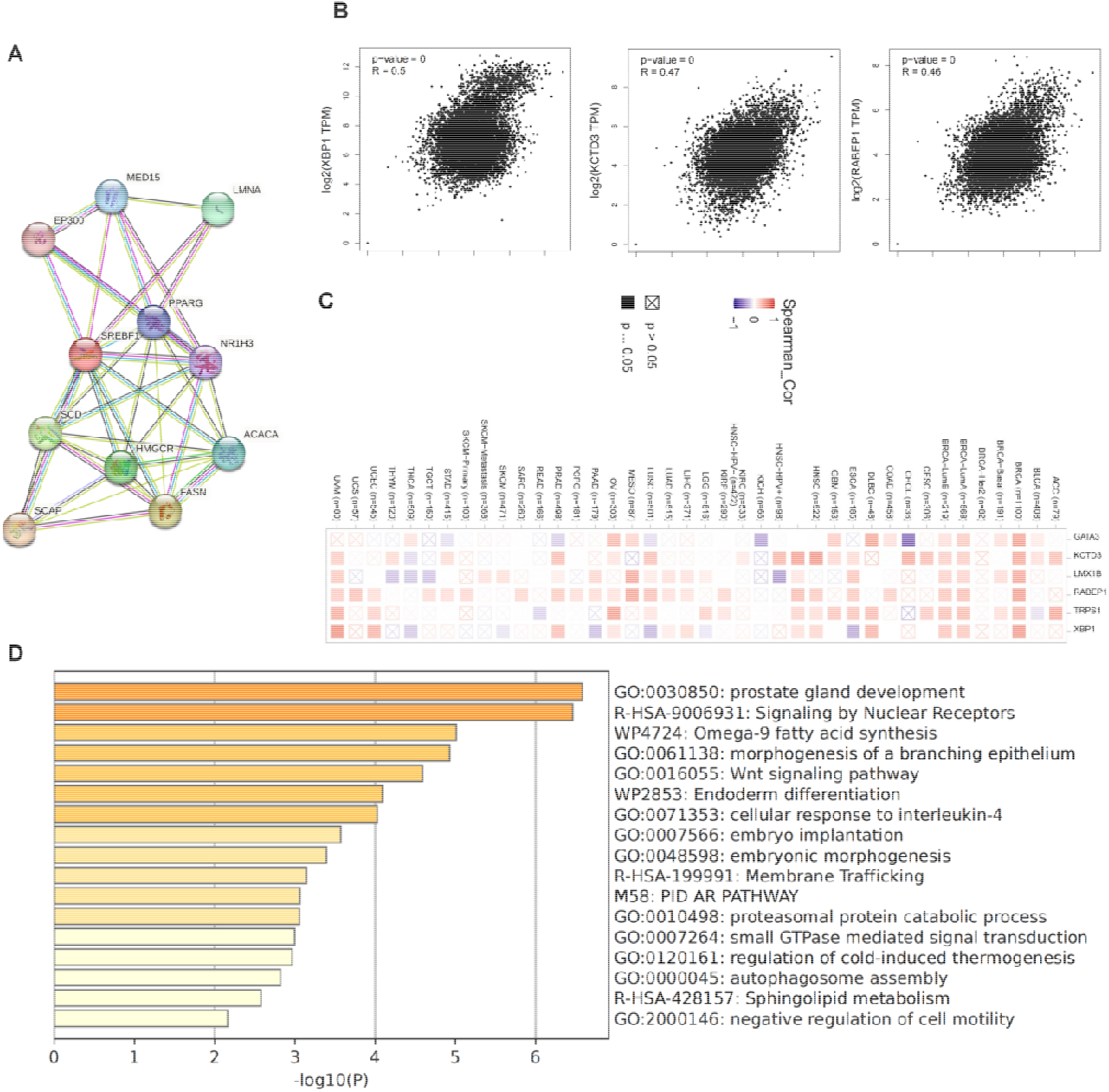
Functional enrichment analysis of SREBF1-related genes. (A) SREBF1-related genes were obtained from the BioGRID web tool, and 15 proteins were displayed. (B) GEPIA2.0 showed the positive correlations between LIPT1 and six genes (TSGA10, ZNF14, EPC2, OSGEPL1, CRBN, and WDSUB1). p-value < 0.001. (C) The heatmap confirmed that SREBF1 expression was positively correlated with the six genes (TSGA10, ZNF14, EPC2, OSGEPL1,CRBN, and WDSUB1) in pan-cancer. (D) GO and KEGG enrichment analyses of SREBF1-related pathway.

### 7. SREBF1 knockdown inhibit cell proliferation and migration

We verified the expression of SREBF1 in human colorectal cell lines, including both normal and tumor cells. Compared to the normal colorectal cell line NCM460, we identified a significantly increase in both mRNA and protein levels of SREBF1 in human colorectal cancer cell lines, particularly in the HCT116 and LS174T cell lines (Figure.9A/9B). To further validate the function of SREBF1 in tumor proliferation and migration, we selected the colorectal cancer cell line HCT116 to study the effect of SREBF1 knockdown on cell (Figure.9C/9D). The colony, CCK8, EDU and transwell results indicated SREBF1 knockdown significantly inhibited HCT116 cell proliferation and migration (Figure.9E-9H). The flow cytometry results demonstrated that SREBF1 knockdown could induce cell cycle arrest, with a decrease in G1 phase and an increase in S phase (Figure.9J). Moreover, SREBF1 knockdown could significantly induce cell apoptosis (Figure.9K/9L).

**FIGURE 9.**
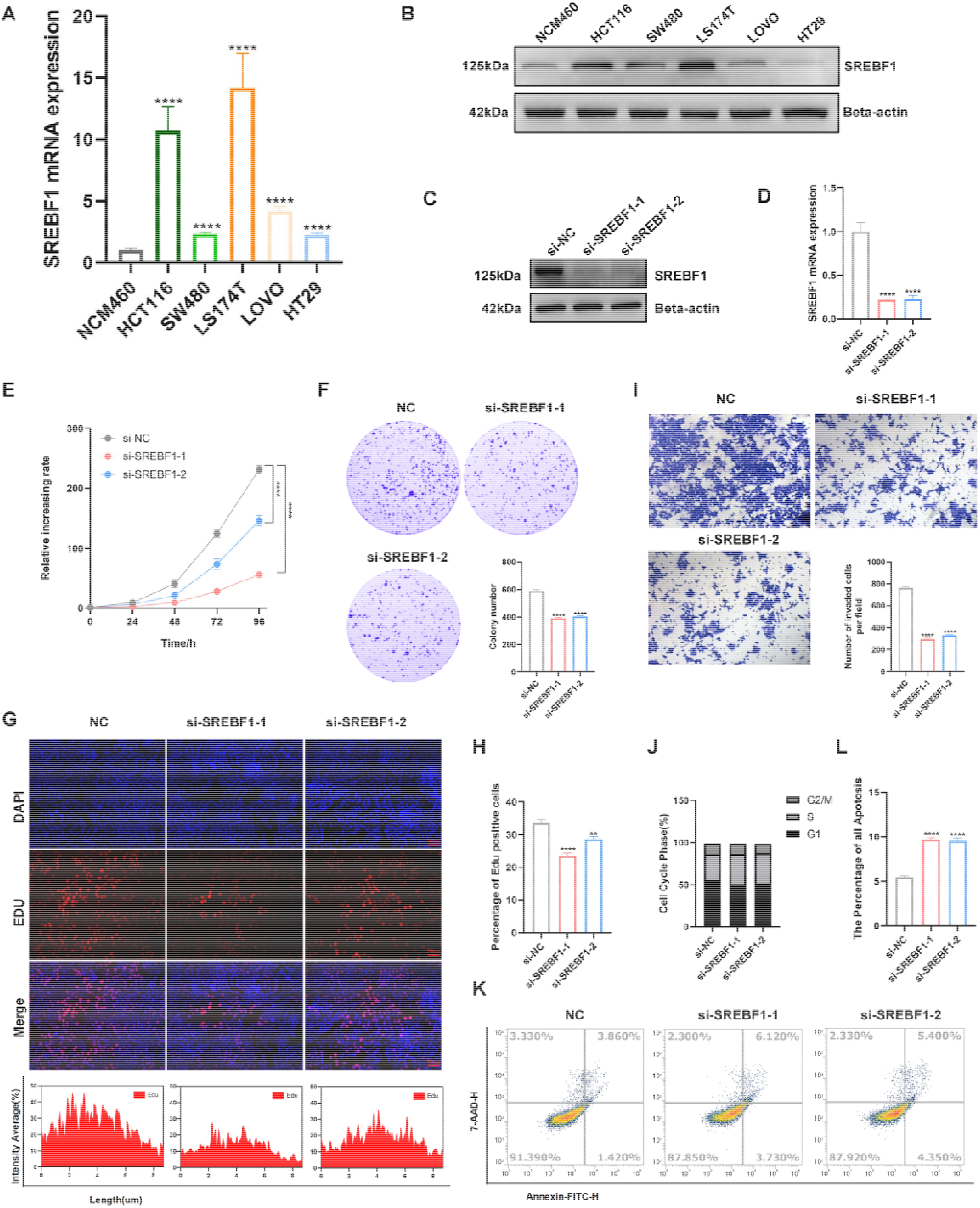
SREBF1 knockdown inhibits the proliferation and migration of HCT116 cell. (A)RT-PCR result of SREBF1 expression in human colorectal normal and tumor cell lines. (B) Western blotting of SREBF1 differential expression in human colorectal normal and tumor cell lines. (C) Western blotting of SREBF1 knockdown in protein level. (D) RT-PCR shows SREBF1 knockdown in mRNA level. (E) Cell proliferation assay by CCK8. (F) Colony assay. (G-H) EDU assay and statistcal anaysis. (I)Transwell assay. (J) Cell cycle analysis by FlowCytometry. ** means p < 0.01;*** means p< 0.001,**** means p< 0.0001.

## Discussion

In earlier studies, SREBF-1 was shown to play a key role in lipid and glucose metabolism, resulting in changes in cell growth, survival and invasiveness. In our research, We comprehensively analysed SREBF1 in several cancers in our research. The expression of SREBF1 was significantly increased in most types of cancers, only in GBM and PCPG with lower expression levels. Patients with high expression level of SREBF1 have been found to have a better prognosis, such as SKCM, BLCA, BRCA, PAAD, UCEC, and STAD. Moreover, we discovered the evident impact of SREBF1 expression on the patients’ stages in CHOL, DLBC, KIRA, and THCA. In addition, we found that SREBF1 had differential expression in weight in different tumors, and speculated that it may be related to cholesterol metabolism, and further studies are still needed to elucidate the specific mechanism.

Mature SREBF1 primarily participates in fatty acid and triglyceride metabolism[22]. In various human cancers, signaling pathway is activated, then promotes the transcriptional activation of SREBP1, thereby facilitating intracellular lipid synthesis in tumor cells[23]. In our study, we found that the expression of SREBF1 varies significantly across some tumors with weight differences. For example, in LUAD and UCEC, the expression of SREBF1 is markedly higher in normal-weight individuals compared to other weight groups (Supplementary Figure.s3). Hence, we thouht that the observed association between SREBF1 expression and patient weight is likely attributable to two aspects: its involvement in metabolic activity and its contribution to immune-related inflammatory responses.

Although few studies have demonstrated that the overexpression of SREBF1 is closely related to many types of tumors, in HCC, the overexpression of SREBF1 is considered to be one of the major factors causing the proliferation and metastasis of cancer cells[24]. In breast cancer, high expression levels of SREBF1 are closely related to adverse outcomes and chemotherapy resistance, its specific mechanism is still not fully understood and requires further research[25]. Based on our research results, we have found SREBF1 genetic alterations, such as mutations and amplification in different forms of cancers. Moreover, there are significant differences in the level of SREBF1 methylation between normal tissues and tumor tissues. The single cell sequence results suggested that the SREBF1-correlated gene might regulate tumor cell biological functions, such as angiogenesis and differential. In addition, combined with GO and KEGG enrichment, several identified biological terms were greatly enriched, indicating processes that are relevant to ‘prostate gland development’, ‘signaling by nuclear receptors’, and ‘omega-9 fatty acid synthesis.

Infiltrating immune cells impact pan-cancer outcomes via dynamic interactions with the tumor microenvironment, influencing tumor progression, immune evasion, and therapeutic response[26]. The study also found a correlation between the SREBF1 expression and plenty of immune infiltration cells in cancer, including CD8+ T cells, Tregs, and myeloid dendritic cells, as well as the immune checkpoints. Currently, our understanding of the functions of SREBF1 in the human immune system is limited. There exists a research gap regarding the roles of SREBF1 in the tumor immune microenvironment, and our findings indicate that focusing on SREBF1 could be a hopeful strategy for immunotherapy. However, the detailed association still needs further investigation, and the role of SREBF1 in TME is worth deep research.

## Conclusion

We systematically analyzed the characteristics of the SREBF1 from many aspects of pan-cancer, including expression, survival, prognosis, genetic mutation, and immune cell infiltration. And further clarified the gene function in colorectal cancer cell growth and migration. The findings indicated that SREBF1 has the potential to be a new prognostic and immune-related biomarker for cancer patients. The study sheds light on the various roles of SREBF1 in different types of cancer and reveals new perspectives on the potential influence of the SREBF1 in pan-cancer.

## Statements and Declarations

### Findings

This work was supported by the National Natural Science Foundation of China under Grant 82003046, the Natural Science Foundation of Chongqing Municipal Science and Technology Bureau under Grant CSTB2022NSCQ-MSX0561, the Chongqing Municipal Health Commission under Grant 2024GDRC011, the Chongqing University under Grant 2023CDJYGRH-YB10, the National Science Foundation of Fujian Province Project under Grant 2020J05212, and the Chongqing Key Laboratory of Emergency Medicine under Grant 2022KFKT01.

## Competing Interests

The authors have no relevant financial or non-financial interests to disclose.

## Author Contributions

All authors contributed to the study conception and design. Material preparation, data collection and analysis were performed by Qinrui Cai, Ling Lin, Xiaoya Zhou, Li Li, Yao Chen, Tianlin Feng, Yuanxiu Gan, Chenhua Zhang and Fan Xu. The first draft of the manuscript was written by Dongling Li and all authors commented on previous versions of the manuscript. All authors read and approved the final manuscript.

## Data Availability

All data included in this study are available by contacting the corresponding authors.

## Notes

### Competing Interest Statement

The authors have declared no competing interest.

