## Supplement Figure for "A comprehensive analysis of the potential biological functions and prognostic values of SREBF1 for multiple cancer types including colorectal cancer"

A

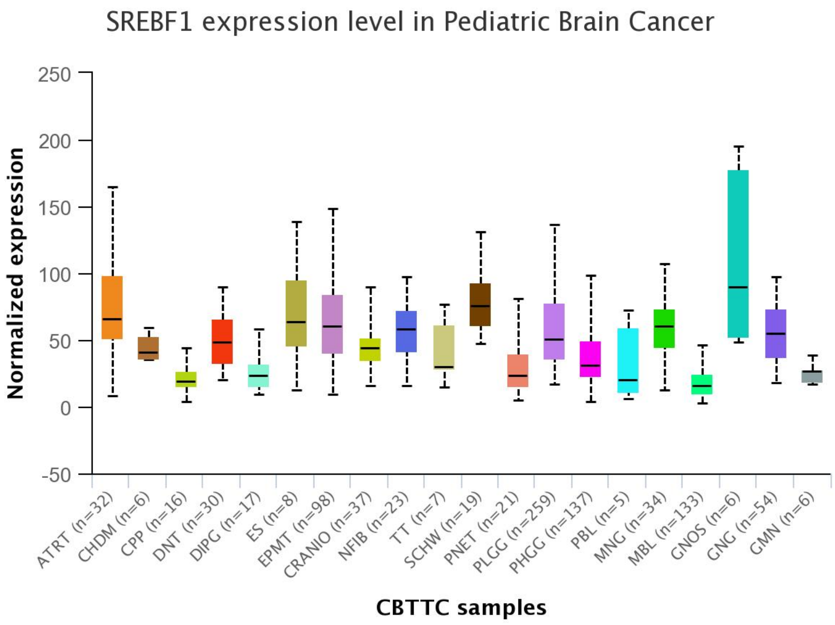


B

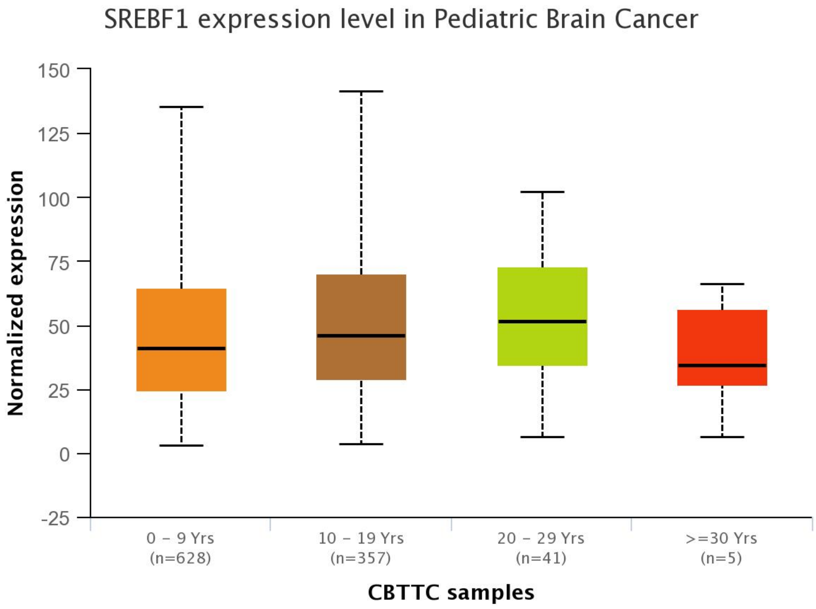


Supplementary Fig.1
(A) SREBF1 expression level in Pediatric Cancer. (B) SREBF1 expression level in Pediatric Cancer of different years.

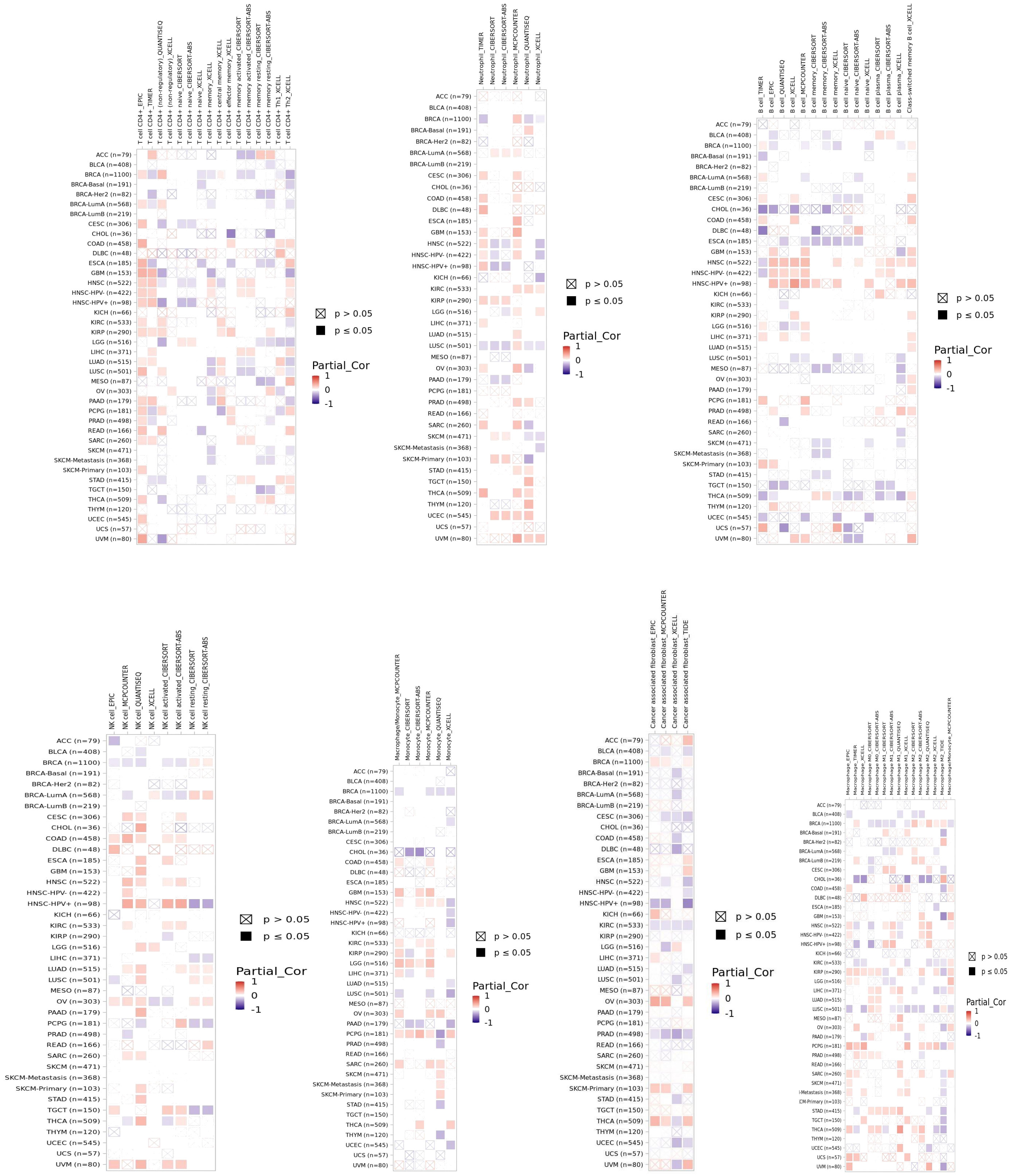


Supplementary Figure 2
The correlation between SREBF1 and the infiltration of natural killer cells (NK), neutrophils, CD4+ T cells, B cells, cancer-associated fibroblasts, and macrophages.

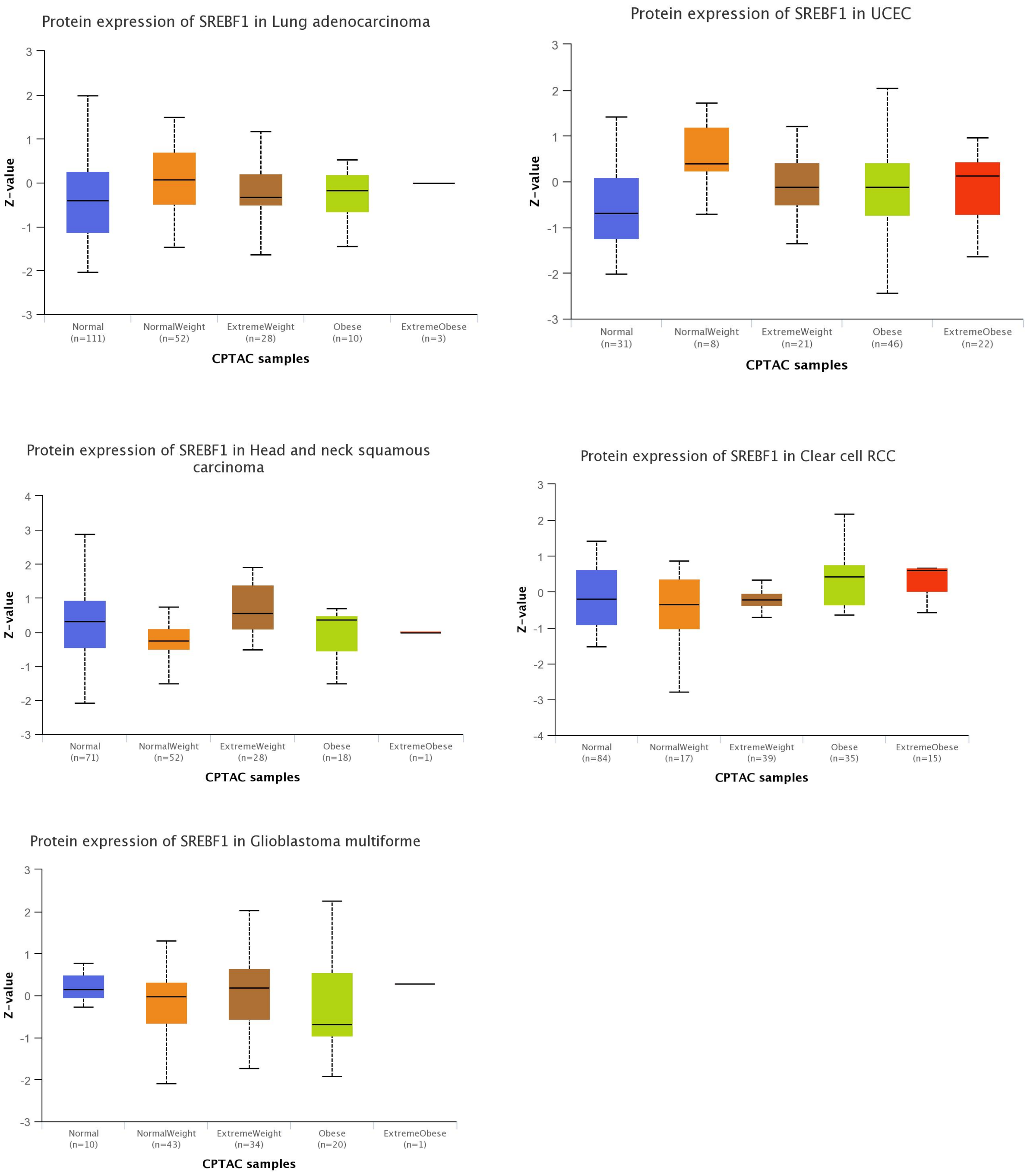


Supplementary Figure 3

SREBF1 expresssion level of preotein in Lung adenocarcinoma, UCEC, Head and neck squamous carcinoma, Clear cell RCC and Gliobastoma  with different weight.
